# DeepSRFusion: a point cloud deep learning framework for super-resolution particle fusion

**DOI:** 10.64898/2026.02.26.708405

**Authors:** Yuyuan Qiao, Jiarui Wang, Jiayi Xi, Jixiang Ding, Tailong Chen, Yiqing Zhang, Lirong Qiu, Weiqian Zhao, Jianli Liu, Fan Xu

**Affiliations:** School of Optics and Photonics, Beijing Institute of Technology, Beijing 100081, China; School of Medical Engineering, Beijing Institute of Technology, Zhuhai 519088, China; School of Medical Technology, Beijing Institute of Technology, Beijing 100081, China; Advanced Research Institute of Multidisciplinary Science, Beijing Institute of Technology, Beijing 100081, China

**Author notes:** These authors contributed equally: Yuyuan Qiao, Jiarui Wang, Jiayi Xi. Correspondence to: Jianli Liu,; Fan Xu.

## Abstract

Deciphering the spatial organization of macromolecular complexes in their native context is central to structural biology. Particle fusion in single-molecule localization microscopy offers a unique capability for high-resolution structural reconstruction *in situ*. However, existing methods face significant challenges from large rotational perturbations and sparse labeling, resulting in compromised accuracy and substantial computational cost. We present DeepSRFusion, a self-supervised pretraining framework for three-dimensional super-resolution particle fusion. By representing single-molecule point clouds as Gaussian Mixture Models, DeepSRFusion integrates data-driven feature learning with physical imaging constraints. A two-stage optimization strategy with dynamic template updating enhances robustness, and a novel Clustering Error metric quantifies fusion quality. Nanometer-scale validation on both simulated and experimental datasets demonstrates high reconstruction fidelity and structural consistency with cryo-electron microscopy and AlphaFold3. DeepSRFusion remains effective under challenging imaging conditions, including large 3D rotations, sparse labeling, high localization uncertainty, and limited particle numbers, while achieving over 100-fold speedups compared to current methods. It resolves fine structural features with a measured spatial resolution of 1.60 ± 0.10 nm, sufficient to distinguish ~10 nm spaced protein pairs and visualize tilted internal substructures within macromolecular assemblies. DeepSRFusion provides a powerful tool for high-precision structural analysis in native cellular environments.

## Introduction

Spatial resolution, the key to elucidating structural biology, has long been constrained by the diffraction limit of optical microscopy**^1,2^**. Single-molecule localization microscopy (SMLM) overcomes this barrier through exploiting the stochastic activation and precise localization of individual fluorescent molecules**^3^**. By sequentially activating sparse subsets of molecules and pinpointing their positions with nanometer precision, SMLM reconstructs super-resolved images that reveal biological structures at unprecedented resolution**^4^**. However, localization uncertainty (LU) caused by photon noise and optical aberrations, as well as insufficient labeling density due to low labeling efficiency, limit the achievable resolution in structural biology studies**^5–7^**.

Particle fusion, a concept originally from the single particle analysis method in cryo-electron microscopy, is a computational approach that enhances the signal-to-noise ratio and reconstruction resolution by merging multiple identical biological structures into an average representation**^8–10^**. In this process, subcellular or macromolecular structures in SMLM, referred to as particles, are segmented from super-resolution images, aligned, and averaged based on structural similarity to produce a two- (2D) or three-dimensional (3D) high-resolution reconstruction**^11^**. Some studies further classify 2D particle images by projection orientation and apply projection reconstruction theory to obtain 3D structures**^12,13^**. Although individual particles suffer from information loss due to limited resolution and labeling density, averaging multiple particles can recover more structural information and achieve sub-10 nm resolution**^14^**. However, SMLM data are fundamentally represented as lists of molecular coordinates with associated uncertainties, rather than pixelated images**^15^**. As a result, the inconsistency between image-based methods and coordinate-based representations can lead to inaccuracies in particle fusion**^16,17^**.

Point cloud analysis plays a central role in computer vision and biological research, enabling the interpretation of spatially resolved molecular and cellular information**^18,19^**. Traditional approaches, such as coordinate-based correlation, incorporate localization uncertainties through statistical pattern recognition and point-set distance functions, thereby improving the accuracy of structural alignment**^16^**. Data-driven strategies, including template-free redundancy analysis and maximum-likelihood model fitting, further enhance alignment robustness and enable inference of macromolecular architectures**^20–22^**. However, traditional coordinate-based methods are highly sensitive to high LU and uneven point densities, often failing under low labeling density (DOL) scenarios**^16^**. While data-driven strategies reduce template bias, they are prone to local optima during optimization, especially for datasets with large spatial orientation variability, and suffer from poor computational efficiency when processing large-scale particle sets due to excessive pairwise alignment**^21,22^**. Additionally, most existing methods lack adaptability across diverse imaging modalities. With the increasing scale of datasets and the complexity of application scenarios, deep learning-based point cloud analysis has emerged as a transformative tool, offering significant improvements in accuracy, efficiency, and robustness**^18,23^**. Recent advances have introduced attention mechanisms, graph matching, and global feature alignment to enhance registration performance and speed**^24,25^**. Furthermore, probabilistic models that integrate Gaussian Mixture Models (GMMs) and Transformer architectures have proven highly effective in handling noise, outliers, and uneven point densities**^26–28^**. Integrating point cloud analysis with particle fusion has the potential to enhance the throughput and accuracy of molecular structure reconstruction**^18^**.

Here we present DeepSRFusion, a point cloud-based <u>deep</u> learning computational framework for <u>s</u>uper-resolution particle <u>fusion</u> in SMLM data. Our method fully accounts for the inherent localization uncertainties and blinking characteristics of fluorescent molecules in SMLM imaging. We convert the point cloud from single-molecule localizations into GMMs to capture the spatial distribution of particles. This representation enables a self-supervised loss that accounts for localization uncertainty and supports robust training of particle fusion network. The specially designed two-stage optimization framework avoids convergence to local optima and produces high-fidelity super-particle reconstructions. Our dynamic template updating strategy significantly reduces reliance on the initial model during iterative optimization. We further developed a novel quantitative metric, Clustering Error (CE), to characterize the error distribution in the super particles formed by particle fusion.

DeepSRFusion effectively harnesses the power of deep learning for feature extraction and successfully integrates it with the inherent physical constraints of SMLM imaging. Nanometer-scale structural validation on both simulated and experimental nuclear pore complex (NPC) datasets demonstrates high reconstruction fidelity and shows structural agreement with cryo-electron microscopy data and AlphaFold3 models. The pretrained framework, incorporating GMMs and tailored optimization strategies, exhibits strong capability under extreme rotational perturbations and low labeling densities, while providing over 100-fold speedups compared to current state-of-the-art methods. DeepSRFusion achieves a spatial resolution of 1.60 ± 0.10 nm for tilted substructures and a spatial distribution uncertainty of 2.59 ± 1.20 nm for individual Nup96 subunits, enabling the discrimination of ~10 nm spaced protein pairs and conformational tilts within macromolecular assemblies. It accurately reconstructs the eight-fold symmetric double-ring architecture of the NPC and resolves tilted sub-modules at specific binding sites. Structural parameters such as ring radius, inter-ring distance, and angular deviation closely match AlphaFold3 predictions. These capabilities enable high-precision *in situ* analysis of macromolecular assemblies and facilitate structural studies in native cellular contexts.

## Results

### DeepSRFusion

DeepSRFusion is a self-supervised deep learning framework specifically designed for 3D particle fusion of SMLM data. The core concept of this framework originates from the inherent nature of SMLM data: due to localization uncertainty (LU), localizations tend to cluster around labeling sites, forming discrete groups**^29^**. This distribution pattern is well represented by a Gaussian Mixture Model (GMM)**^27^**(**Supplementary Note 1.1**). To capture this spatial structure, we designed a deep point cloud network that models single-molecule localizations as GMMs, enabling accurate analysis and globally consistent particle registration (**Fig. 1a, Extended Data Fig. 1**). We further designed a self-supervised loss function tailored to the characteristics of SMLM data, which incorporates LU by modeling each point as a Gaussian distribution (**Supplementary Note 1.2**). This formulation enables probabilistic comparison between point clouds, jointly accounting for spatial coordinates and localization errors. By optimizing the alignment of these Gaussian-modeled representations, the proposed method significantly improves the stability and robustness of registration, particularly in highly symmetric macromolecular assemblies with large rotational perturbations.

**Fig. 1:**
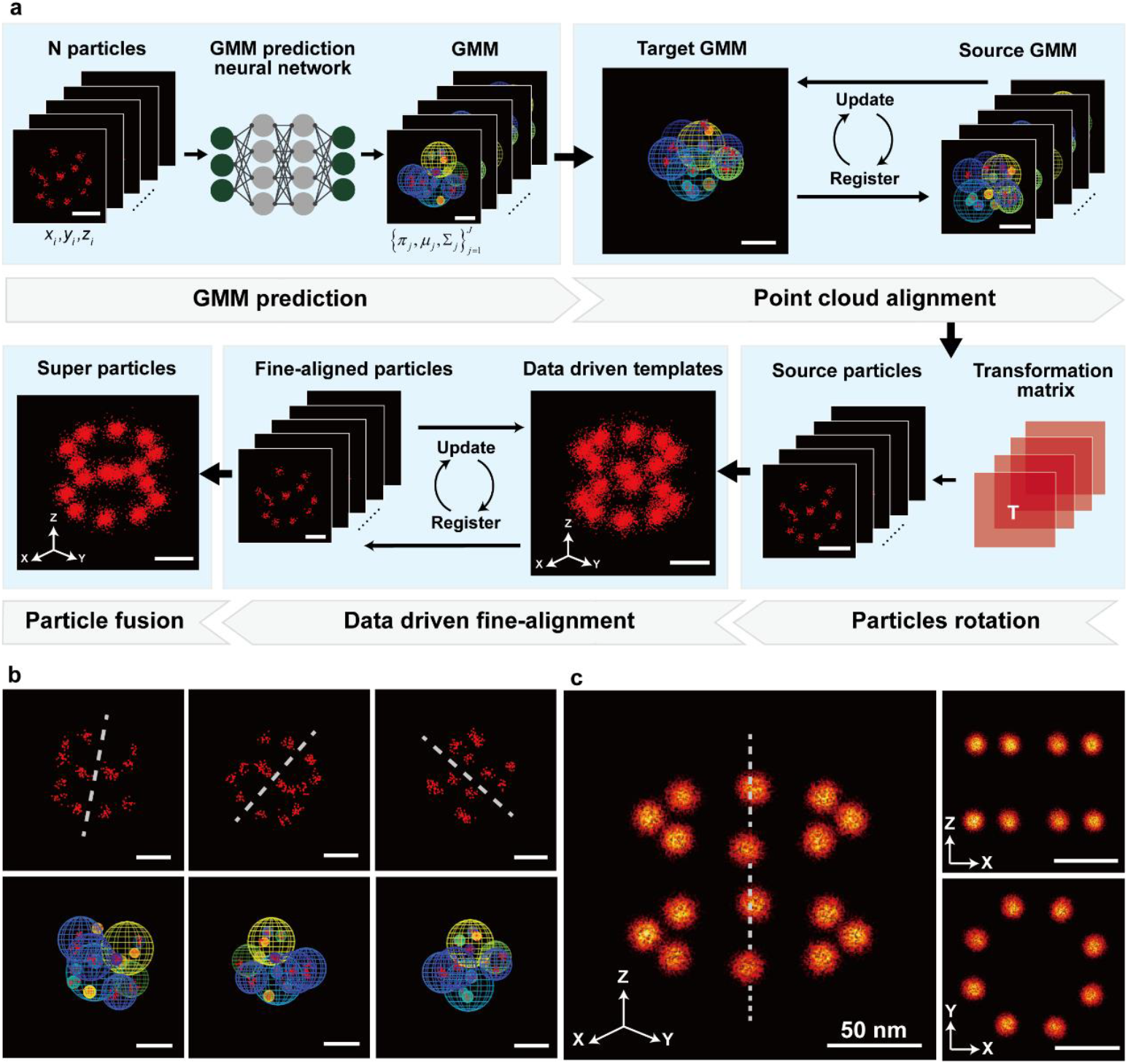
DeepSRFusion concept. **a**, the workflow begins with a pre-trained point cloud network generating GMM representations for input 3D particles, each containing localization uncertainties. During alignment, the algorithm randomly selects one particle’s GMM as the initial target template, then registers all remaining source GMMs to this target to compute rigid transformations. Through iterative processing, transformed source GMMs are superimposed onto the dynamically updated target until convergence is achieved, yielding a GMM model, which then computes the transformation matrix T for point cloud alignment. This matrix subsequently aligns all particle point clouds, which fuse into a data-driven template. The pipeline concludes with fine registration cycles that progressively refine particle-template alignment, ultimately generating the final high-precision “super particle” reconstruction. **b**, GMM predictions from the alignment network, demonstrating consistent modeling of particle point clouds across diverse 3D orientations. **c**, 3D, side, and top views of particle fusion results from DeepSRFusion (100 particles). Dashed lines indicate particle central axes. Data were simulated using Nup107 structures with 70% density of labeling (DOL), 4 nm localization uncertainty (LU), and random rotations (±180° along *xyz*-axes). Bars were 50 nm.

The overall workflow of DeepSRFusion is illustrated in **Fig. 1a**. The algorithm takes as input a set of *N* particles, where each particle is defined as *P*={*p*_1_,*p*_2_,…,*p*_i_,…,*p*_n_}, comprising *n* localized 3D points. Each point p =(*x*_*i*_,*y*_*i*_,*z*_*i*_,Σ_*i*_) includes its spatial coordinates (*x*_*i*_,*y*_*i*_,*z*_*i*_)∈R^3^ and a corresponding LU Σ_*i*_, which reflects the inherent imprecision in SMLM data. To model the spatial distribution of each particle, a pre-trained deep neural network is employed to map the input localizations to a GMM (**Supplementary Note 1.3**). This results in a set of predicted GMMs, G={*G*_1_,*G*_2_,…,*G*_*k*_,…,*G*_*N*_}, representing the ensemble of input particles. Each GMM *G*_*k*_ is defined as a weighted sum of Gaussian components:

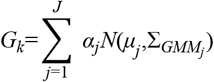

where *α*_*j*_ is the mixture weight, and *N*(*μ*_*j*_,Σ_*j*_) denotes a Gaussian distribution with mean *μ*_*j*_ and covariance matrix 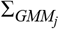.

A specially designed GMM-based neural network was dedicated to point cloud registration (**Extended Data Fig. 1**). Rather than directly aligning point clouds, DeepSRFusion performs end-to-end optimization under the expectation-maximization framework, where the network predicts GMM components that guide the alignment process. To minimize the influence of the initial model on the alignment, we proposed a dynamic template updating strategy embedded in the iterative optimization procedure (**Extended Data Fig. 2**). The process designates a randomly selected particle’s GMM as the target template, with the others as sources. Iterative optimization then estimates rigid transformation of each source to the target, aggregates the aligned sources to update the target, and outputs the final transformations, thereby reducing initial bias.

A two-stage optimization framework is employed to avoid convergence to local optima and to produce high-fidelity super-particle reconstructions. The first stage uses a GMM-based neural network for point cloud alignment. In the second stage, a data-driven fine alignment is conducted to enhance registration accuracy through iterative optimization (**Supplementary Note 1.4**). An initial template is constructed by aggregating all input particles, serving as the spatial reference for fine registration. During each iteration, for every source particle, the target point cloud is created by excluding this source particle from the current template to avoid self-bias. The source particle is then registered to this target by optimizing rigid transformation parameters, guided by a Gaussian distance metrics that integrates SMLM-specific localization uncertainty. After registering all source particles in one iteration, the data-driven template is updated with these registered particles to incorporate refined spatial information. This “register-update” cycle repeats until convergence, yielding accurately aligned particles for high-resolution structural reconstruction.

The pre-trained DeepSRFusion network exhibits strong generalization, enabling direct application to particles with diverse imaging parameters, including varying localization uncertainties and labeling densities. Validated on the NPC dataset (**Fig. 1b, c**), the model accurately reconstructs 3D structures across orientations and effectively handles large initial rotations and structural symmetry, unlocking new capabilities for the structural analysis of biological complexes.

### Validation and Performance Benchmarking of DeepSRFusion Framework

To assess reconstruction accuracy, we began by validating the DeepSRFusion framework using simulated Nup107 data under controlled conditions (**Supplementary Note 2**). In SMLM, uncertainty in molecule positions leads to clustering around labeling sites, forming discrete groups**^30^**. To quantitatively assess reconstruction accuracy, we introduced a clustering-based error metric, Clustering Error (CE), which exploits the intrinsic property that both individual particles and their fused super-particles naturally form Gaussian-distributed clusters around labeling sites (**Fig. 2a, Supplementary Note 5.1**). CE provides a statistical measure of alignment quality by assessing the compactness of localization clusters, enabling rigorous evaluation of reconstruction performance across diverse biological scenarios.

**Fig. 2:**
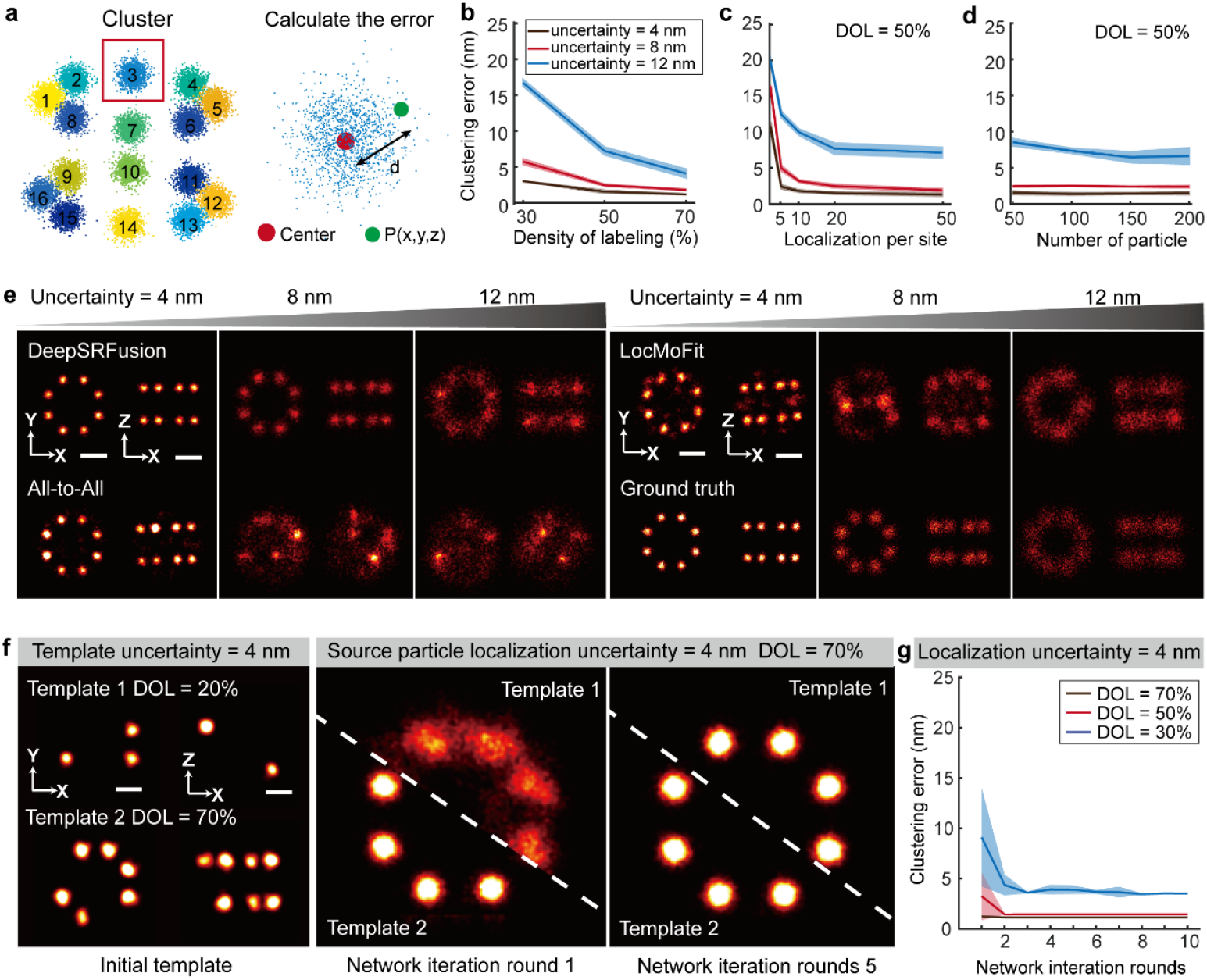
Validation of DeepSRFusion using simulated Nup107 datasets. **a**, Clustering error calculation based on Euclidean distances from k-means cluster centroids (k = 16). **b-d**, Final results after fine alignment. **b**, Reconstruction performance across gradients of labeling efficiency (DOL = 30%, 50%, 70%) and localization uncertainty (LU = 4 nm, 8 nm, 12 nm) within a rotation range of ±180°. **c**, Effect of localization per site (LPS = 5, 10, 20, 50) under DOL = 50%. For each parameter setting, 100 particles were used for reconstruction, and the experiments were repeated five times with independent datasets. **d**, Consistency analysis with varying particle numbers (50-200 particles) under DOL = 50%. **e**, Comparative reconstruction quality (±180° rotations; DOL = 50%, LPS = 5, 100 particles). **f**, Template bias elimination tested with low-quality (template 1, DOL = 20%, 3 sites) and high-quality (template 2, DOL = 70%, 11 sites) initial templates. The comparison of the reconstruction results of network iteration 1 and 5 under the uncertainty of source particle localization = 4 nm and DOL=70% is shown on the right. **g**, Quantitative template bias analysis (initial templates: DOL = 20-70%; source particles: DOL = 30-70%). Error bands indicate the standard deviations of the measured data.

We evaluated the impact of different density of labeling (DOL) and LU on particle fusion performance. Specifically, datasets were generated with DOL levels of 30%, 50%, and 70%, and LU values of 4 nm, 8 nm, and 12 nm, within a rotation range of ±180° (**Fig. 2b**). Under optimal conditions (DOL = 70%, LU = 4 nm), DeepSRFusion achieved high accuracy (CE = 1.15 ± 0.02 nm). At 30% DOL, CE remain acceptable with LU = 4 nm and 8 nm (3.00 ± 0.02 nm and 5.71 ± 0.36 nm, respectively). When LU increased to 12 nm, CE rose significantly, indicating reduced reconstruction quality due to lower data fidelity. This trend persists across all tested rotation angles and was observed consistently in both GMM network and data-driven fine alignment stages (**Supplementary Fig. 1**).

We investigated the influence of LPS (Localizations Per Site) on reconstruction quality under a fixed DOL. For DeepSRFusion, both the CE from GMM-based neural network predictions and the refined outputs from the data-driven processing remained stable when LPS ≥ 5. Specifically, at an average of 5 LPS, the CE values of the final reconstructions were 2.32 ± 0.27 nm, 4.91 ± 0.36 nm, and 12.43 ± 0.43 nm for LUs of 4 nm, 8 nm, and 12 nm, respectively. In comparison, at 50 LPS, the corresponding CE values decreased to 1.23 ± 0.18 nm, 1.84 ± 0.16 nm, and 7.10 ± 0.70 nm (**Fig. 2c, Supplementary Fig. 2a, b**). These results demonstrate the robustness of DeepSRFusion under sparse localization conditions. Across varying LU levels, the reconstruction quality remained consistent and was insensitive to particle number under 50% DOL and ±180° rotation constraints. Furthermore, while the reconstruction quality continued to improve with increasing particle count at LU = 12 nm, it remained stable at LU = 4 nm and 8 nm, results obtained with 50 particles were comparable to those with 200 particles. This indicates that, under high-quality data conditions, DeepSRFusion can achieve high-fidelity reconstructions even with a limited number of particles (**Fig. 2d, Supplementary Fig. 2c, d**).

Furthermore, we conducted comparative benchmarking against established methods**^21,22^** (All-to-All and LocMoFit) under realistic imaging conditions (DOL = 50%, LPS = 5, LU = 4 nm, 8 nm, 12 nm) within rotation range of ± 180 ° and ± 15 ° (**Fig. 2e, Extended Data Fig. 3, Supplementary Fig. 3, Supplementary Note 4.1**). While all methods performed comparably for mild rotation (±15°), DeepSRFusion maintained superior accuracy under extreme orientation (±180°), achieving a CE of 2.32 ± 0.27 nm at LU = 4 nm, whereas conventional methods showed substantial degradation. Visual inspection at LU= 4 nm confirmed DeepSRFusion preserved architectural features without the hotspot artifacts (All-to-All) or the misalignment errors (LocMoFit) observed in other methods. As data quality decreased (LU = 8, 12 nm), the performance of All-to-All and LocMoFit deteriorated significantly, while DeepSRFusion remained stable and closely matched the ground truth. In addition, under different localization uncertainties and particle number conditions, DeepSRFusion requires less processing time than the other two methods, with speedup ratios ranging from 52 to 185 (**Extended Data Fig. 3, Supplementary Fig. 3**). This result indicates that DeepSRFusion substantially enhances the time efficiency of data analysis while maintaining clustering accuracy, making it well suited for large-scale, high uncertainty single-molecule localization datasets.

Finally, we examined the performance of DeepSRFusion in mitigating template bias through GMM-based iterative optimization (**Fig. 2f**). Remarkably, the method remained effective even under extreme conditions (DOL = 20%, LU = 4 nm, 3 detectable sites), surpassing typical experimental requirements. Across templates with 20–70% DOL, reconstructions consistently converged within five iterations despite initial template variations (**Extended Data Fig. 2**). As the number of iterations increases, the CE values across all conditions exhibited convergence, and the final standard deviation error significantly decreases. After 10 iterations, reconstructions at DOL = 30%, 50%, and 70% showed minimal variation (**Fig. 2g, Extended Data Fig. 4**). These results establish DeepSRFusion as a robust solution for particle fusion, particularly excelling in challenging orientations and template bias mitigation.

### Particle fusion of 3D SMLM data with rotational robustness

Biological structures are inherently distributed in 3D space with diverse orientations**^31^**. Traditional fusion methods are typically designed for small-angle, uniform datasets**^21,22^**. In contrast, DeepSRFusion accommodates arbitrary rotation angles, enabling robust alignment across a wide range of orientations. To evaluate its performance under extreme rotational variability, we applied it to Nup96**^32,33^**, a core component of the nuclear pore complex (NPC) that forms octameric subunits with eight-fold symmetry across cytoplasmic and nuclear rings, totaling 32 copies per NPC (**Fig. 3a**). Using networks pre-trained on simulated data, we applied GMM prediction directly to Nup96 STORM data (**Fig. 3b, Supplementary Fig. 4a, Supplementary Note 2**). The network successfully recovered the characteristic double-ring architecture of NPCs in both *xy*- and *xz*-views. These results demonstrate the strong generalization capability of DeepSRFusion and its robustness to unseen conditions, providing a reliable foundation for high-fidelity 3D particle fusion.

**Fig. 3:**
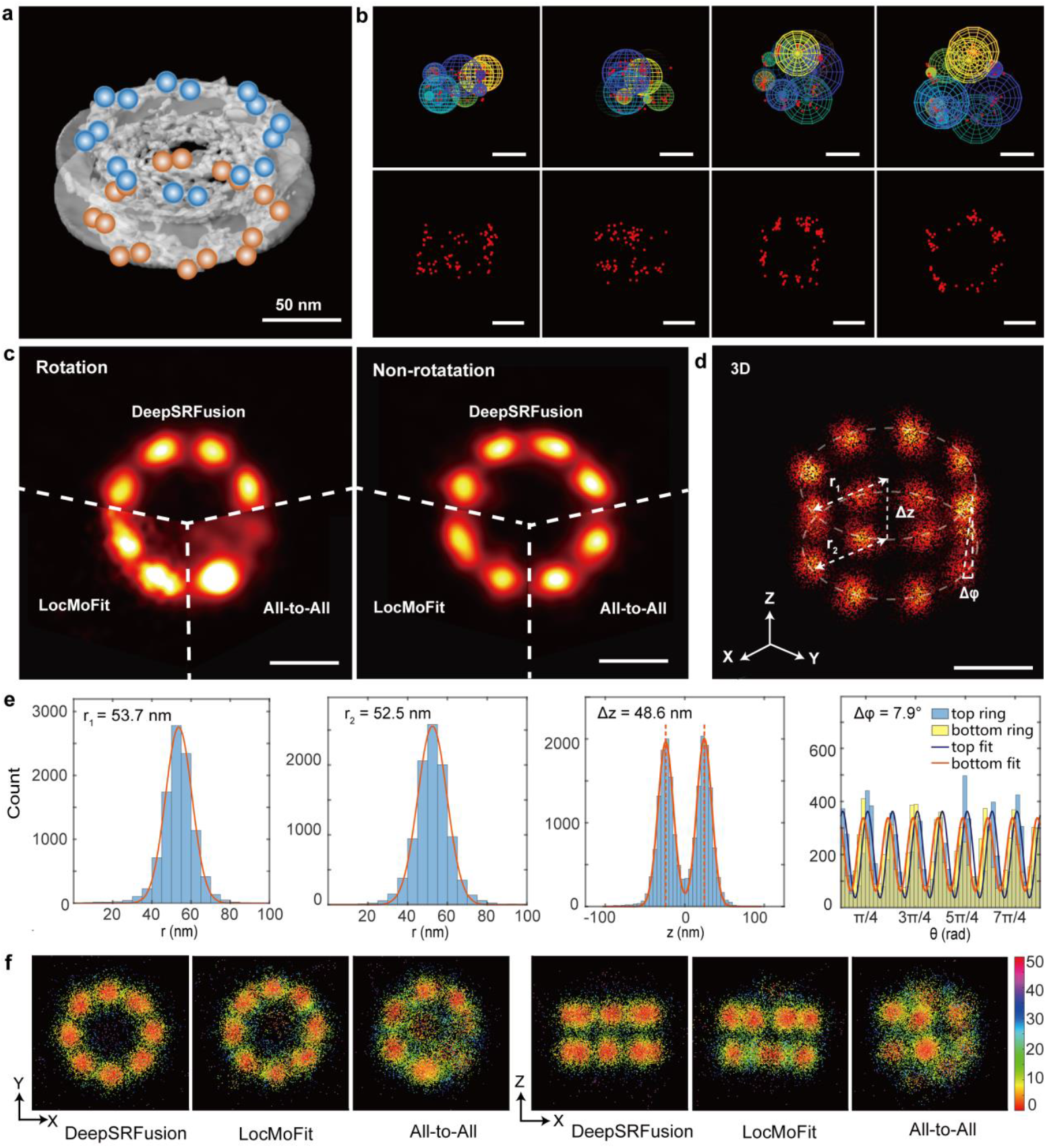
Validation of DeepSRFusion based on Nup96 protein datasets. **a**, Cryo-EM structure of Nup96 showing two concentric 16-site rings (https://www.ebi.ac.uk/emdb/). Blue particle pairs form top ring of NPC; orange particle pairs form bottom ring of NPC. **b**, Representative single-particle STORM image (upper row) and corresponding GMM prediction (bottom row). **c**, Reconstructed Nup96 structures from ±180°rotated particles (left) and non-rotated controls (right) using DeepSRFusion, LocMoFit, and All-to-All (n=200 particles). **d**, 3D reconstruction from 200 rotated STORM particles. **e**, Localization histograms quantifying structural parameters under extreme rotation angle: ring radii (upper/lower), inter-ring distance, and angular deviation. **f**, Clustering error (k=16) across labeled sites, DeepSRFusion illustrating no hotspot artifacts (mis-localization clustering seen in LocMoFit or All-to-All) and clear site differentiation. Bars were 50 nm.

We evaluated reconstruction performance of three methods (DeepSRFusion, All-to-All, LocMoFit) under two conditions: a rotation group (±180° random rotations about *xyz*-axes) and a non-rotation group (original orientations) (**Fig. 3c**). In the non-rotation group, all methods successfully reconstructed the double-ring, eight-fold symmetric NPC structure. However, under large rotational variance, performance diverged markedly. DeepSRFusion maintained high-quality reconstructions with dense, well-defined clusters (**Fig. 3d, Supplementary Fig. 4b**), whereas All-to-All exhibited severe hotspot artifacts and failed to preserve structural integrity, consistent with simulation results. LocMoFit preserved overall architecture but showed deformation and peripheral noise due to misalignment. These differences were further emphasized by CE visualization, with DeepSRFusion achieving the lowest and most uniform error distribution (**Fig. 3f**). Parallel experiments with Nup107 datasets-a conserved nucleoporin subunit that constitutes the scaffold of the NPC, exhibiting the characteristic double-ring structure of NPCs**^34^** (**Extended Data Fig. 5**) confirmed these trends, demonstrating DeepSRFusion’s superiority under high angular variability while performing comparably to existing methods at low rotations.

To assess reconstruction accuracy, key structural parameters such as inter-ring distance (Δz), ring radii (r_1_, r_2_), and inter-ring angular deviation (Δφ) of Nup96 reconstructions were quantified. For the rotation group, measurements were r_1_ = 53.7 nm, r_2_ = 52.5 nm, Δz = 48.6 nm, and Δφ = 7.9° (**Fig. 3e**). Corresponding values for the non-rotation group were r_1_ = 54.3 nm, r_2_ = 51.8 nm, Δz = 49.0 nm, and Δφ = 8.5° (**Supplementary Fig. 5**). These measurements closely match published EM data**^35,36^**, confirming consistent reconstruction fidelity across initial angular conditions. Notably, the rotation group exhibited a smaller ring radius difference (0.4 nm) compared to the non-rotation group (1.6 nm), reflecting the inherent challenge of resolving near-identical symmetric features under extreme rotations. This symmetry-induced convergence behavior highlights both the robustness and intrinsic limitations of high-symmetry reconstructions.

### Nanometer-scale structural reconstruction of Nup96 subunits

To assess the ability of DeepSRFusion to resolve high-resolution features, we used the RESI (Resolution Enhancement by Sequential Imaging) dataset (**Supplementary Note 4.3**), which offers near Ångström-scale precision sufficient to resolve individual Nup96 subunits. To validate the predicted subunit organization and spatial arrangement, we employed AlphaFold3**^37^** to generate a structural model and estimate the distribution of Nup96 labeling sites (**Fig. 4a, Extended Data Fig. 6, Supplementary Note 5.3**). Consistent with previous studies**^35^**, AlphaFold3 predicted that each Nup96 ring contains eight protein pairs, yielding a total of 32 labeling sites across upper and lower rings. The predicted geometry includes an upper ring radius r_1_ = 54.8 nm, lower ring radius r_2_ = 54.6 nm, inter-ring distance Δz = 52.8 nm (**Fig. 4b**). Additionally, the angular deviation Δφ = 6.1 ± 0.3°, adjacent Nup96 proteins are spaced 11.6 ± 0.1 nm laterally (**Fig. 4c**).

**Fig. 4:**
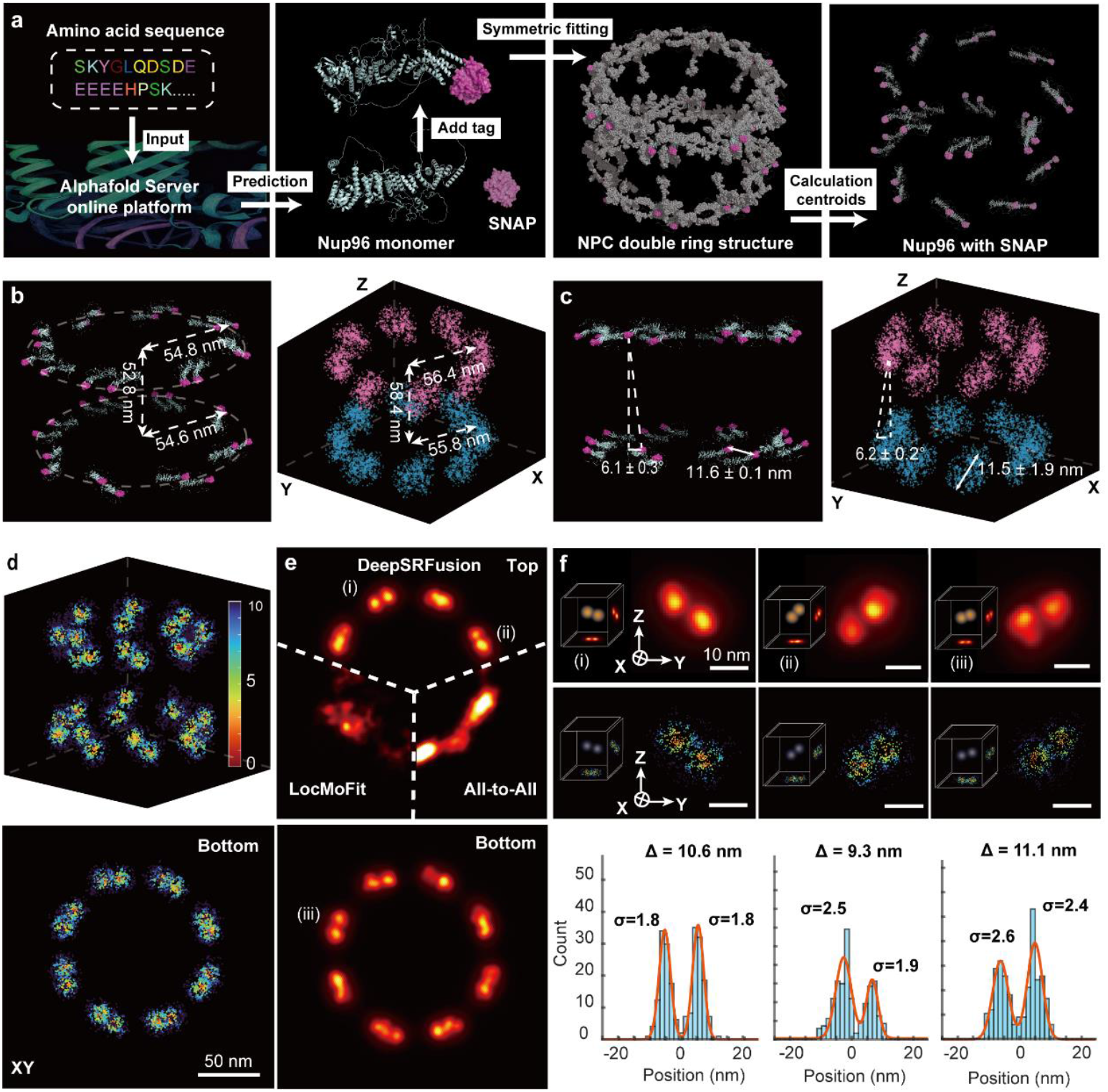
Validation of DeepSRFusion based on RESI Nup96 experimental data. **a**, Alphafold3 prediction Nup96 structure workflow. **b-c**, Alphafold3 (left column) and DeepSRFusion (right column) predicted Nup96 fundamental structure: ring radii (upper/lower), inter-ring distance(b), angular deviation and adjacent Nup96 proteins lateral distance (c). **d**, CE values after particle fusion on the RESI Nup 96 dataset with added random rotation angles in DeepSRFusion. The lower panel shows CE values of bottom ring in *xy*-view. **e**, Particle fusion reconstruction results of DeepSRFusion contrasted with LocMoFit and All-to-All, (i) - (iii) represent three Nup96 protein pairs. **f**, Calculation of localization point distribution of labeled (i) - (iii) Nup96 protein pairs in (e) for distance between two sites, including super-resolution reconstruction map (upper row), error plot (middle row), particle pair lateral distance distribution chart (bottom row).

To further evaluate the robustness of DeepSRFusion under arbitrary orientations, we applied high-angle 3D rotations to the dataset and reconstructed the rotated particles. From RESI-processed data, DeepSRFusion predicted an upper ring radius r_1_ = 56.4 nm, lower ring radius r_2_ = 55.8 nm, inter-ring distance Δz = 58.4 nm (**Fig. 4b)**, and angular deviation Δφ = 6.2 ± 0.2°, adjacent Nup96 proteins are spaced 11.5±1.9 nm laterally (**Fig. 4c**). We applied trigonometric fitting to model the Nup96 subunits’phase shift *θ* in both the cytoplasmic and nucleoplasmic rings, which represents their relative rotational displacement in angular distribution. By comparing *θ*_CR_ and *θ*_NR_, the angular deviation Δφ between the two rings can be quantitatively determined, providing a measure of their spatial misalignment. To account for stochastic estimation errors arising from point cloud measurement noise, angular uncertainty was quantified through a bootstrap analysis. This involved 1,000 bootstrap iterations of random sampling with replacement, each followed by constrained trigonometric fitting. The resulting empirical distribution of Δφ provided a robust measure of angular uncertainty (**Supplementary Fig. 6, Supplementary Fig. 7**). This close agreement confirms the tilted arrangement of Nup96 subunits in native NPC architecture and provides experimental validation for AI-predicted conformations.

To evaluate the robustness of DeepSRFusion under arbitrary orientations, we applied ±180° random rotations along all axes to 1,191 RESI-processed Nup96 particles and performed reconstruction (**Fig. 4d**). Despite the extreme rotational variability, DeepSRFusion accurately resolved the canonical eight-fold symmetric double-ring NPC architecture and further distinguished two tilted sub-modules within individual Nup96 subunits (**Fig. 4e, Extended Data Fig. 7**). These sub-modules correspond to specific binding sites, with a spatial distribution uncertainty of 2.59± 1.20 nm for individual Nup96 subunits (**Extended Data Fig. 8, Extended Data Fig. 9**). To assess the spatial resolution achieved by DeepSRFusion, we performed a Fourier Shell Correlation (FSC) analysis on the reconstructed Nup96 structures. FSC estimates resolution by comparing the similarity between two independently reconstructed 3D volumes across spatial frequencies**^38^**. This analysis yielded a resolution of 1.60 ± 0.10 nm for the two tilted sub-modules within individual Nup96 subunits (**Extended Data Fig. 10**). High structural resolution was maintained even under extreme rotational conditions, demonstrating robust adaptability to diverse imaging scenarios.

Quantitative analysis of y-axis distance distributions for representative protein pairs (**Fig. 4f**) revealed inter-site distances of 10.6 nm, 9.3 nm, and 11.1 nm. Statistical analysis across all pairs yielded a mean spacing of 10.16±1.82 nm (**Extended Data Fig. 9**), with individual measurements deviating by ≤1.5 nm from the mean. These results align closely with both the previously reported value**^37^** and AlphaFold3 predictions, validating accurate resolution of ~10 nm-separated labeled sites. This nanometer-scale measurement capability enables precise structural characterization of macromolecular complexes such as the NPCs, supporting mechanistic insights at the molecular level.

## Discussion

In this study, we introduced DeepSRFusion, a deep learning-based particle fusion framework that integrates GMMs with point cloud registration to address key challenges in SMLM. By reformulating the point cloud alignment as a probabilistic matching task between GMM representations, our method explicitly accounts for spatial uncertainties inherent in localization data. A self-supervised pretraining stage, tailored to SMLM characteristics, enables robust training and accurate registration under challenging conditions such as large rotational variations (±180°) and low labeling density. Comprehensive validation using both simulated and experimental datasets demonstrates that DeepSRFusion consistently achieves high-fidelity reconstructions, closely matching structural benchmarks from Cryo-EM and AlphaFold3 models. These results confirm the method’s capability to resolve nanometer-scale features such as protein pairs spaced at ~10 nm and tilted conformations within macromolecular assemblies, marking a significant advance in SMLM-based structural biology.

A central innovation of DeepSRFusion lies in the integration of a pretrained GMM-based neural network to probabilistically represent SMLM point clouds. This representation captures localization uncertainties while offering a robust and natural metric for structural comparison and alignment. By circumventing direct point-to-point distance calculations, which are highly sensitive to noise and outliers, our method achieves superior performance on symmetric and heterogeneous assemblies. The two-stage optimization strategy, coupled with dynamic template updating, effectively mitigates initial model bias and avoids convergence to suboptimal local minima, ensuring accurate and biologically consistent reconstructions. Additionally, the proposed CE metric provides an intuitive and quantitative measure of reconstruction quality, particularly in evaluating the spatial compactness and accuracy of localized clusters under diverse imaging conditions.

Despite these advances, several limitations and avenues for future development remain. Its applicability to more complex biomolecular systems, particularly those characterized by high structural heterogeneity, low symmetry, or dynamic conformational variability, has yet to be fully explored. Extending the framework to a wider range of targets, such as membrane receptors**^39,40^**, cytoskeletal assemblies**^41^**, and transient complexes**^42^**, will be essential for establishing broader generalizability. While the current architecture is computationally feasible, future iterations could adopt more advanced designs to better capture fine structural variations and improve scalability. Additionally, extending the method to incorporate multi-color SMLM data could enhance molecular specificity and enable more comprehensive structural analyses.

In conclusion, DeepSRFusion represents a substantial advancement in computational methodologies for SMLM data analysis, effectively bridging deep learning-based feature extraction with physically informed probabilistic modeling. By enabling robust and high-resolution particle fusion under realistic imaging conditions, the framework provides a powerful tool for *in situ* structural biology, facilitating the quantitative reconstruction of macromolecular architectures within their native cellular environments.

## Code availability

The DeepSRFusion toolbox is developed for three-dimensional particle fusion in single-molecule localization microscopy. It includes a test case with step-by-step instructions that guide users through the complete particle fusion workflow, from model training to fusion of both simulated and experimental datasets. Further updates and additional resources will be continuously made available freely on the Github repository as of the date of publication.

## Data availability

Data underlying the results presented in this paper are not publicly available at this time but may be obtained from the authors upon reasonable request.

## Acknowledgements

We thank Pingyong Xu from Institute of Biophysics, Chinese Academy of Sciences for the valuable suggestions on the manuscript. We thank Jonas Ries and Yu-Le Wu from European Molecular Biology Laboratory for their helpful clarifications regarding data preprocessing, as well as for providing the open-source dataset used in LocMoFit. This work was supported by the National Natural Science Foundation of China (62272041), the National Key R&D Program of China (2023YFC3402600) and Postdoctoral Fellowship Program of CPSF (GZB20250982).

## Extended Data Figure

**Extended Data Fig. 1:**
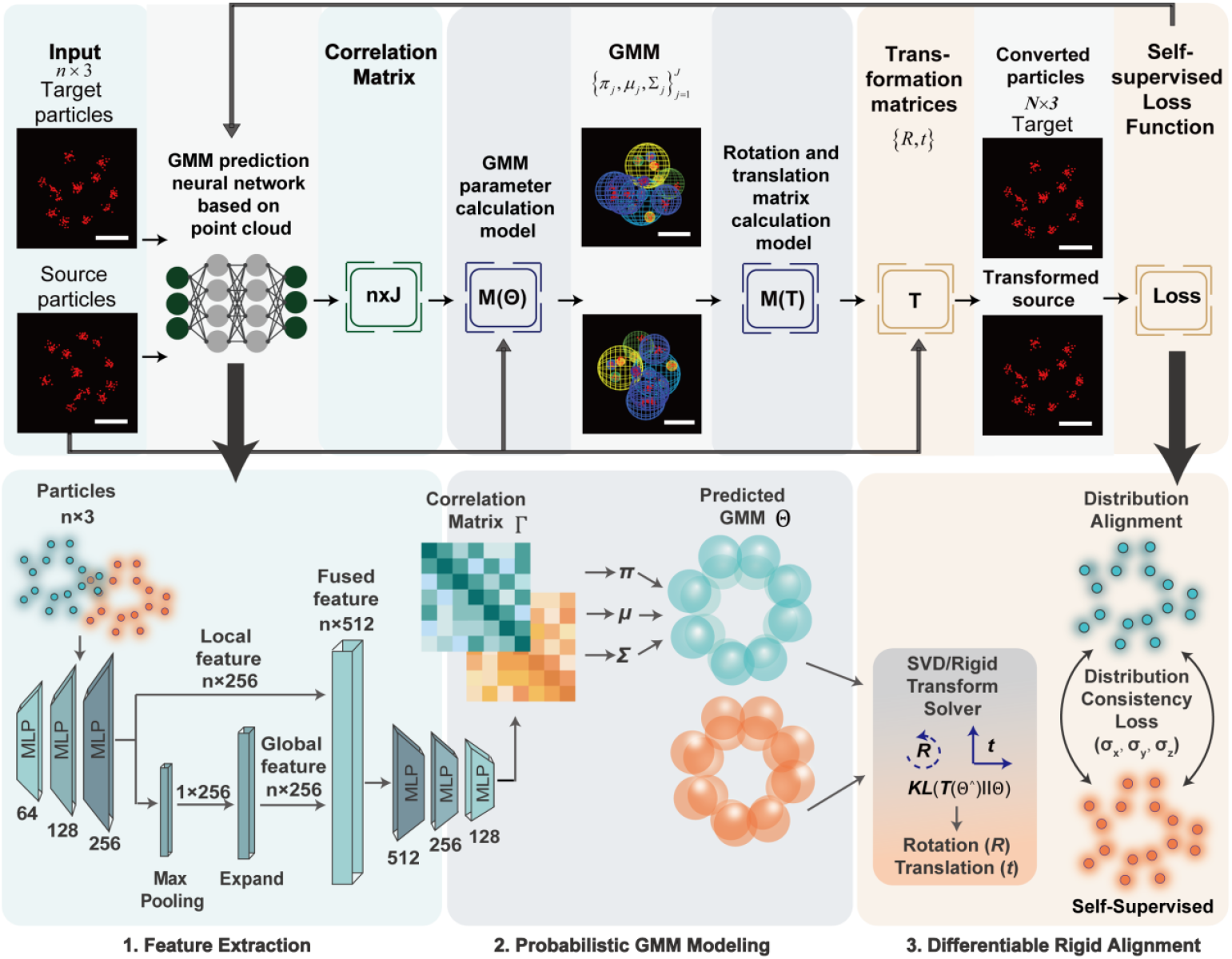
Deep learning point cloud registration network structure of DeepSRFusion. Schematic diagram illustrating the architecture of the GMM-based deep neural network dedicated to point cloud registration in the DeepSRFusion framework. The network adopts a three-module design to achieve end-to-end, probabilistic alignment of SMLM point clouds, while explicitly accounting for inherent LU of SMLM data. Prediction network takes 3D SMLM point clouds, each point defined by spatial coordinates *x, y, z* as input. It employs a PointNet-based backbone consisting of multi-layer perceptrons (MLPs) with hidden layer sizes 64, 128, and 256, to extract local point features. A max-pooling operation generates global structural features, which are concatenated with local features to form fused feature representations (dimension: *n* × 512, where *n* is the number of points in the input cloud). Subsequent MLPs (512, 256, 128) output an *n* × J correlation matrix *Γ* with row sums normalized to 1, where each element *γ*_*ij*_ denotes the latent correspondence probability between the *i*-th input point and the *j*-th Gaussian component of the target GMM. GMM parameter calculation module analytically derives GMM parameters 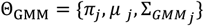 (mixture weights, mean vectors, and covariance matrices of Gaussian components) from the correspondence matrix *Γ* and input point coordinates. It enforces isotropic covariance matrices *Σ*_*j*_ = *diag*([Σ_*j*_ ^2^, *σ*_*j*_ ^2^, *σ*_*j*_ ^2^]) to simplify computation and enhance robustness to noise; variance*σ*_*j*_ ^2^ is calculated via inner-product substitution in the covariance update to avoid overfitting to outliers. Rigid transformation calculation module computes optimal rigid transformation parameters T = (*R*, *t*) (rotation matrix *R* and translation vector *t*) to align the source point cloud to the target. It minimizes the Kullback-Leibler (KL) divergence between the GMM of the transformed source cloud and the target GMM, which is mathematically equivalent to minimizing a weighted Mahalanobis distance loss using the correspondence probabilities *γ*_*ij*_ as weights. The optimal *R* is solved via weighted singular value decomposition (SVD) of a cross-covariance matrix, while *t* is derived by centering the weighted means of source and target GMMs. Notably, this network design replaces the traditional Expectation-Maximization (EM) algorithm’s E-step with learned correspondence prediction, reducing the risk of convergence to local optima and improving alignment robustness for symmetric macromolecular structures (e.g., nuclear pore complexes) under large rotational perturbations (±180°) and low labeling density.

**Extended Data Fig. 2:**
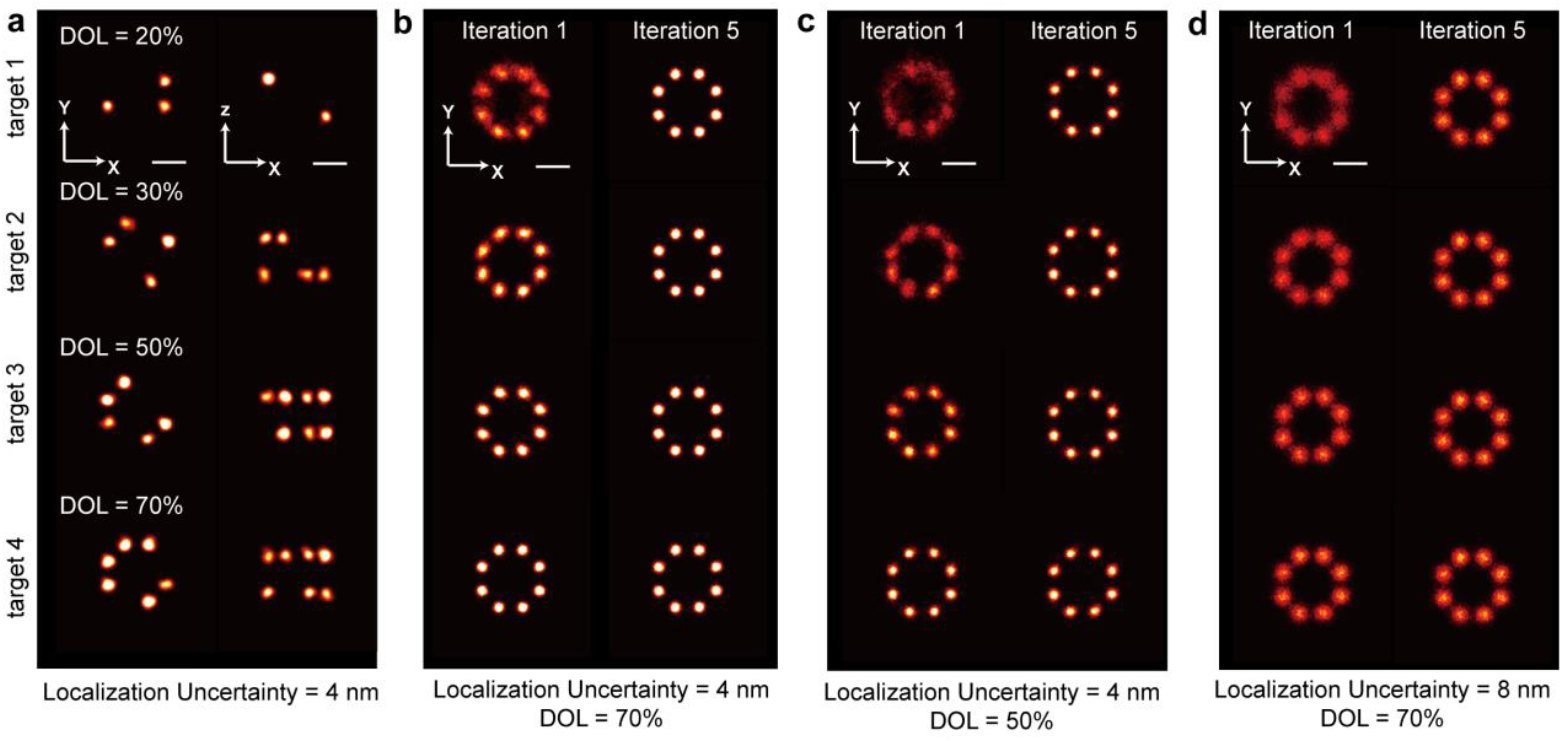
Analysis of the impact of initial templates on reconstruction results. **(a)** Initial template quality under different DOL conditions. Target 1,2,3,4 randomly retain the results generated from 3,5,8,11 valid sites, respectively. **(b-d)** Reconstruction results of particles with different parameters under various initial templates. As the number of iterations increased to 5, the template deviation significantly decreased. The final reconstruction results under all conditions achieved visually consistent good quality. Bars were 50 nm, 100 particles were used for reconstruction.

**Extended Data Fig. 3:**
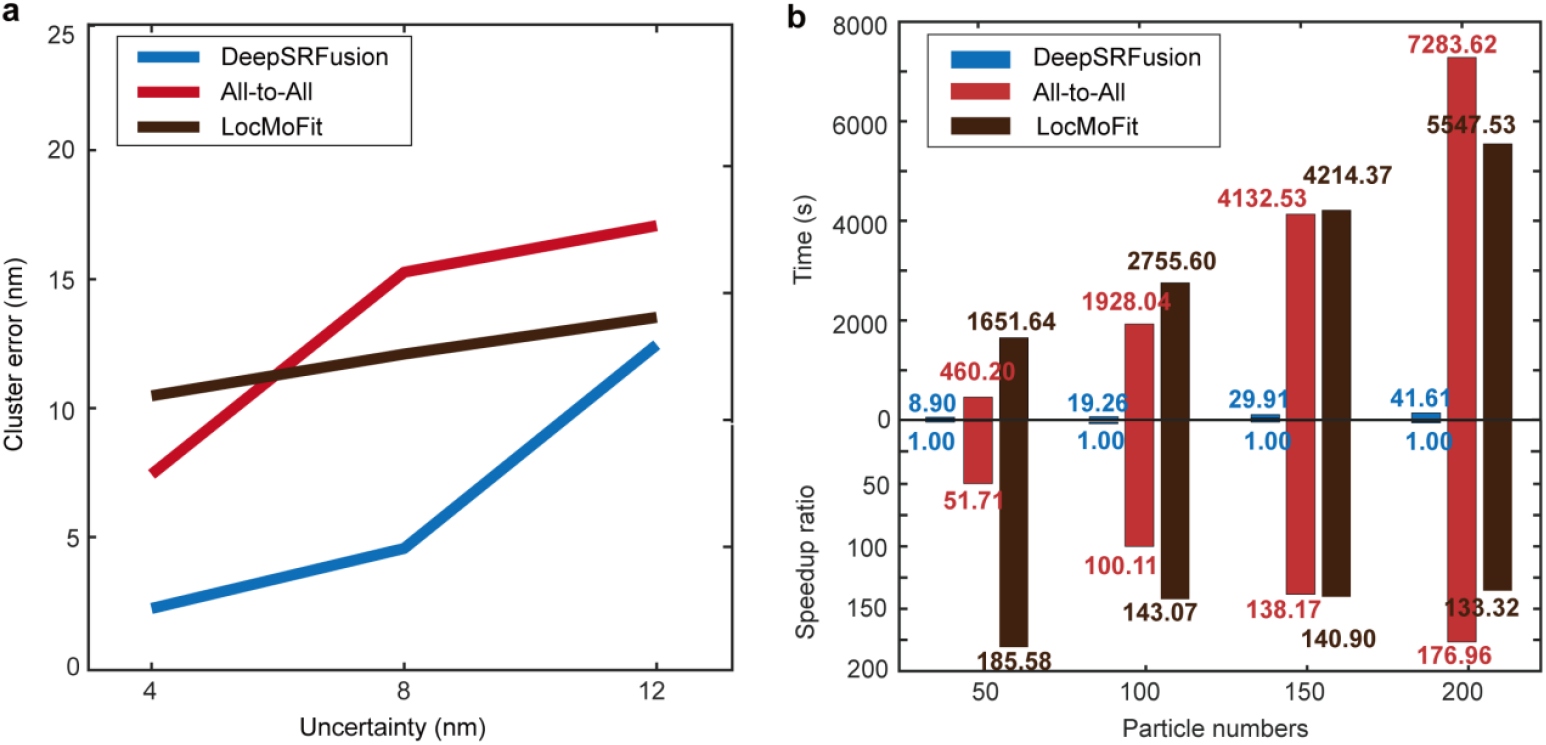
DeepSRFusion, All-to-All, and LocMoFit performance under extreme rotation (± 180°). **(a)** CE statistic under extreme rotation with DOL=50%, LPS=5, 100 particles, and different localization uncertainty. **(b)** Processing time (upper) and speedup ratio comparison (bottom) under extreme rotation with different particle numbers.

**Extended Data Fig. 4:**
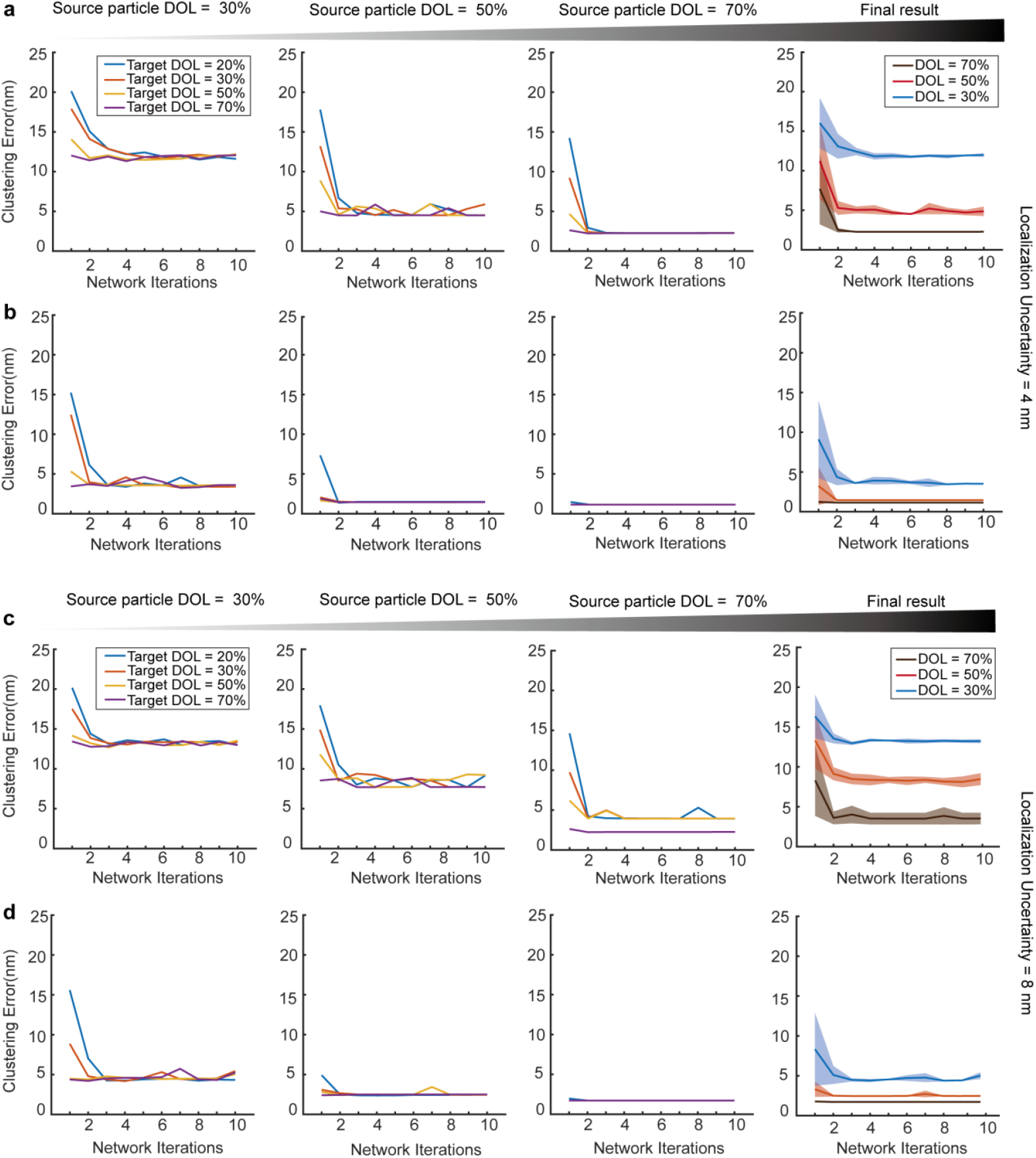
Analysis of the impact of initial template selection on CE in reconstruction results. **(a-b)** CE statistic of GMM-based neural network for point cloud registration and data-driven fine alignment under LU = 4 nm, respectively. Final result indicates the mean CE of iteration process for different DOL datasets under the conditions of each initial template. The shaded area represents the corresponding standard deviation, which is an integrated display of all curves in the left diagram. **(c-d)** CE statistic of GMM-based neural network for point cloud registration and data-driven fine alignment under LU = 8 nm, respectively.

**Extended Data Fig. 5:**
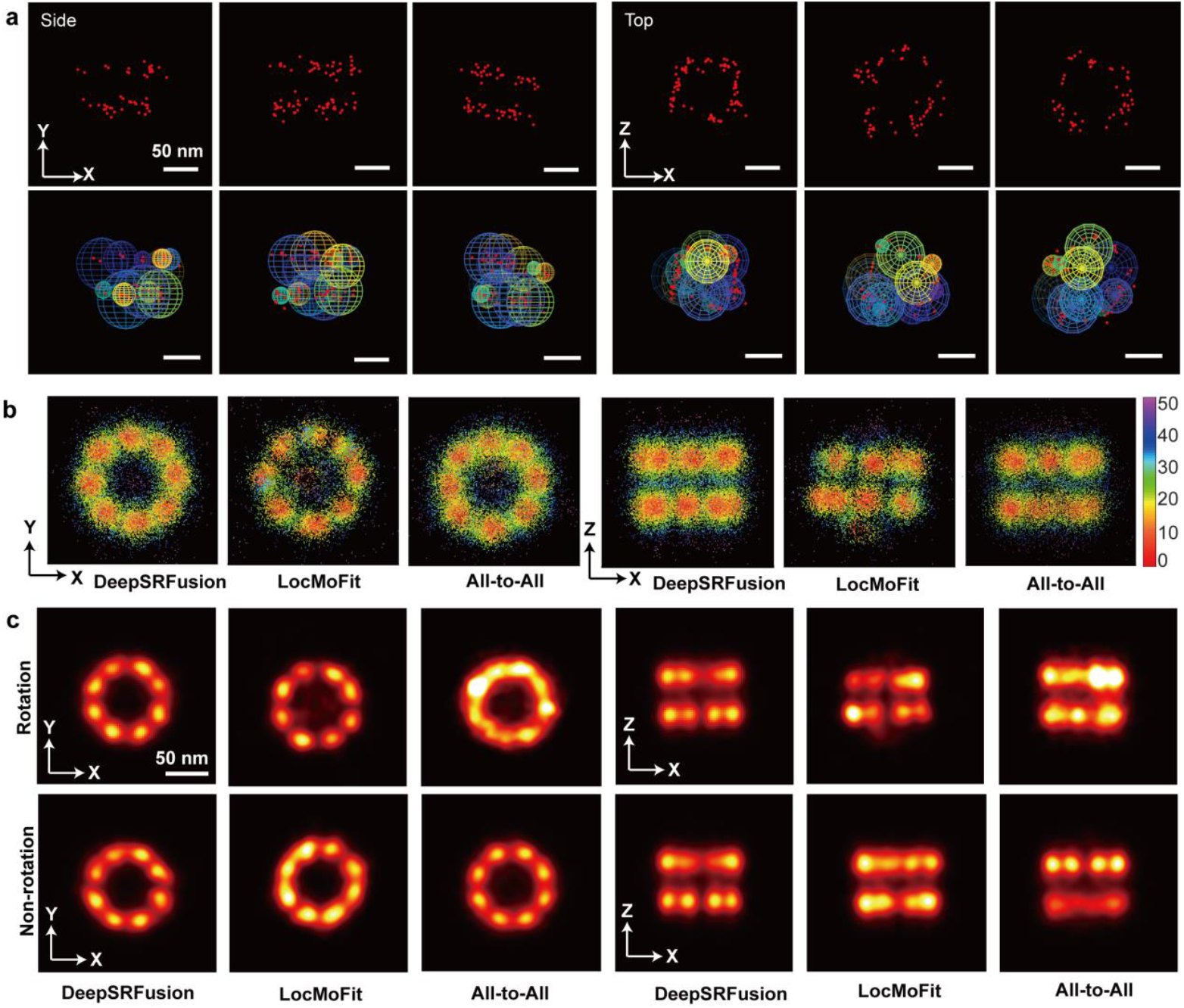
Validation of DeepSRFusion based on Nup107 protein datasets. **(a)** The Nup107 single particle of STORM imaging from side view and top view, and the corresponding GMM predicted by DeepSRFusion. **(b)** Clustering error (k=16) across labeled sites of Nup107, DeepSRFusion illustrating no hotspot artifacts (mislocalization clustering seen in LocMoFit or All-to-All) and clear site differentiation. **(c)** Reconstructed Nup107 structures from ±180°rotated particles (left) and non-rotated controls (right) using DeepSRFusion, LocMoFit, and All-to-All (n=200 particles). Bars were 50 nm.

**Extended Data Fig. 6:**
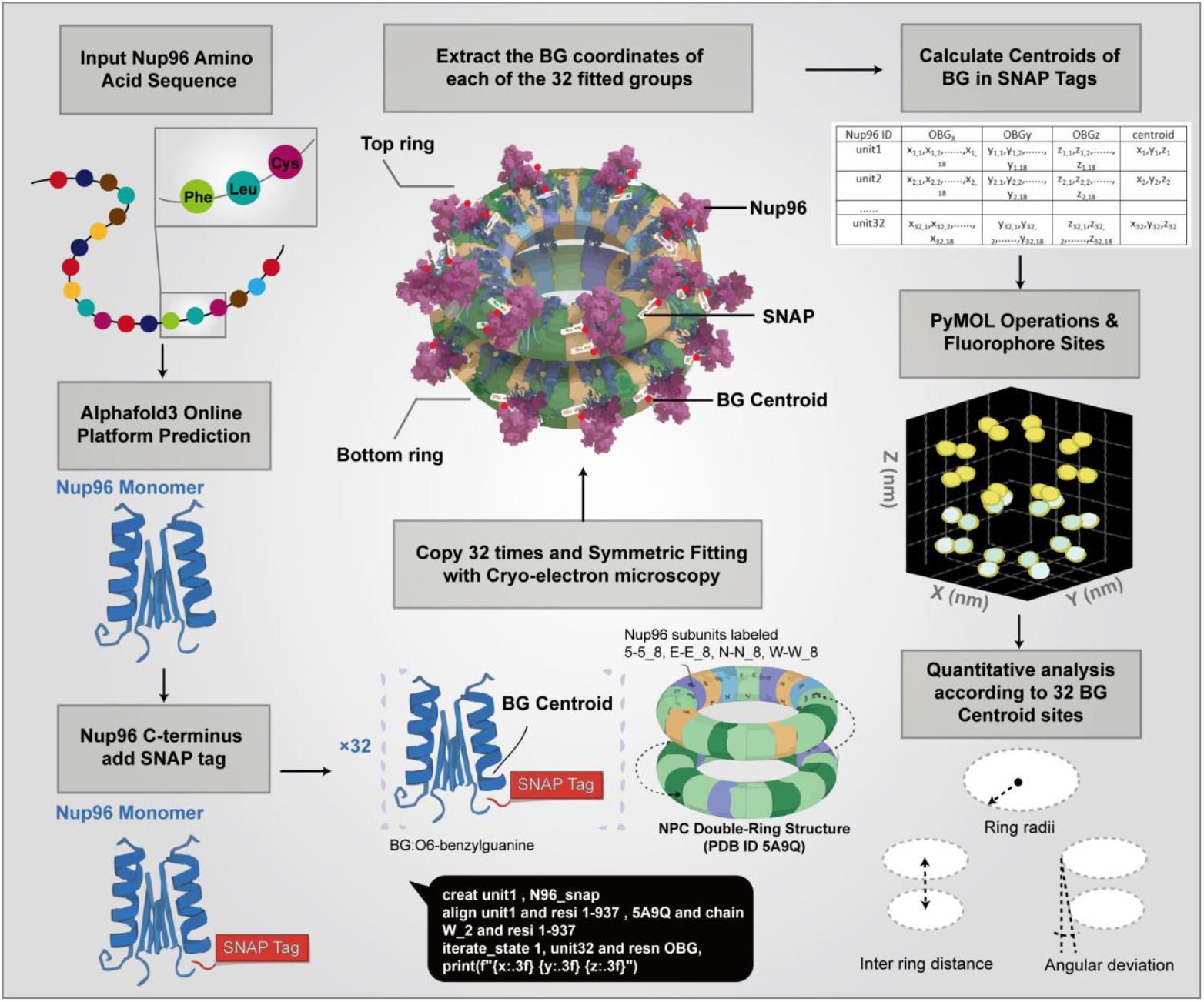
Workflow for predicting and fitting nuclear pore complex (NPC) architecture based on AlphaFold3. The amino acid sequence of Nup96 was used as input for AlphaFold3 to generate a monomeric structural model, to which a SNAP tag was appended at the C-terminus. This tagged monomer was then rigidly fitted to the 32 corresponding Nup96 positions in the cryo-EM NPC structure (PDB 5A9Q). In PyMOL, the OBG (O6-benzylguanine) atoms of all 32 SNAP tags were extracted, converted from Å to nanometer, and used to compute the centroid position of each fluorophore. These 32 centroid coordinates were subsequently used to calculate key geometric parameters of the NPC double-ring architecture, including ring radii, axial spacing, XY-projected distances between paired loci, and the angular relationships between upper and lower subunits (**Supplementary Note 5.3**)

**Extended Data Fig. 7:**
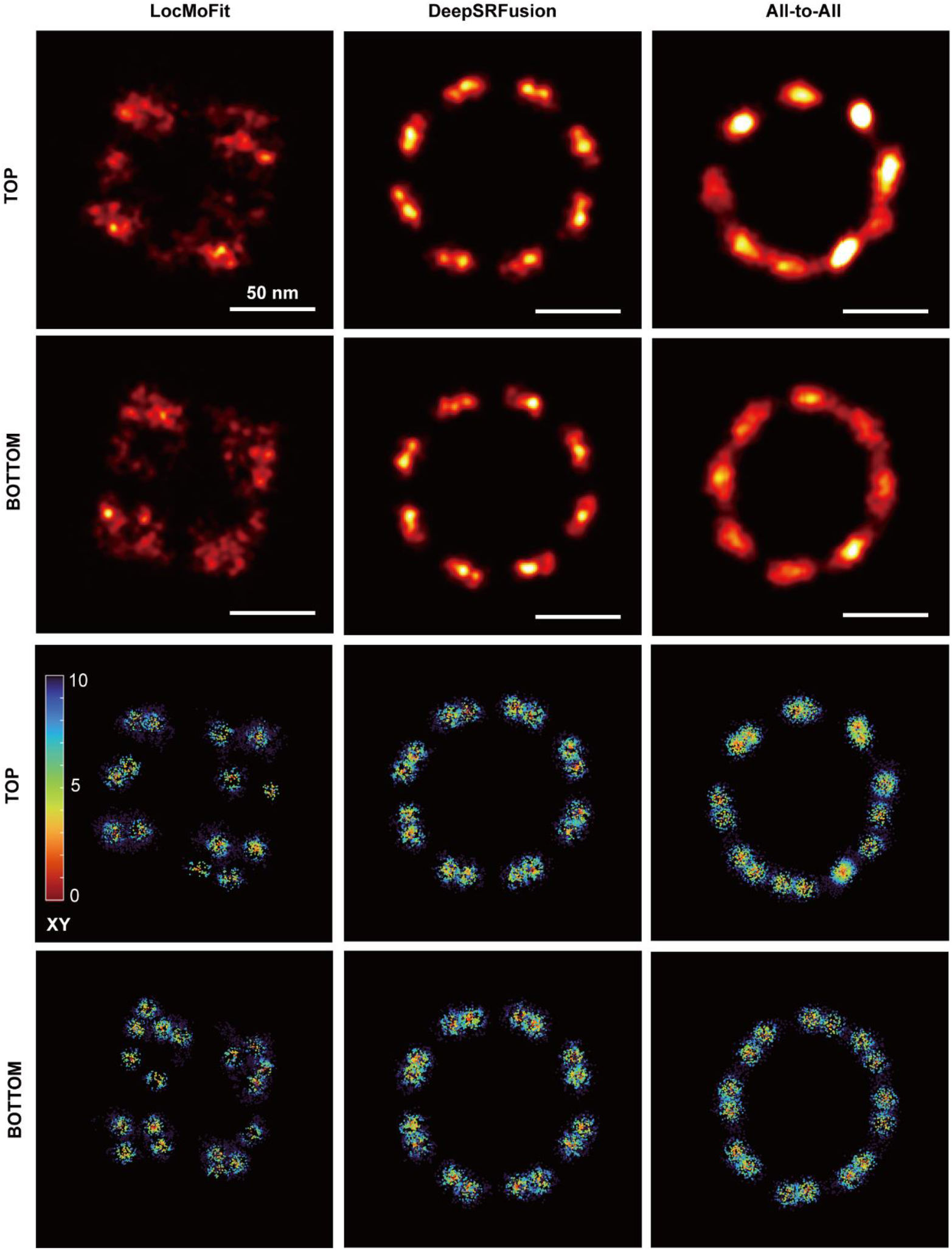
The reconstruction results of RESI Nup96 among three methods. The upper two rows are the reconstructed top/bottom ring structure of Nup96, and the lower two rows are the error plot of corresponding reconstructed ring under CE analysis. Bars were 50 nm.

**Extended Data Fig. 8:**
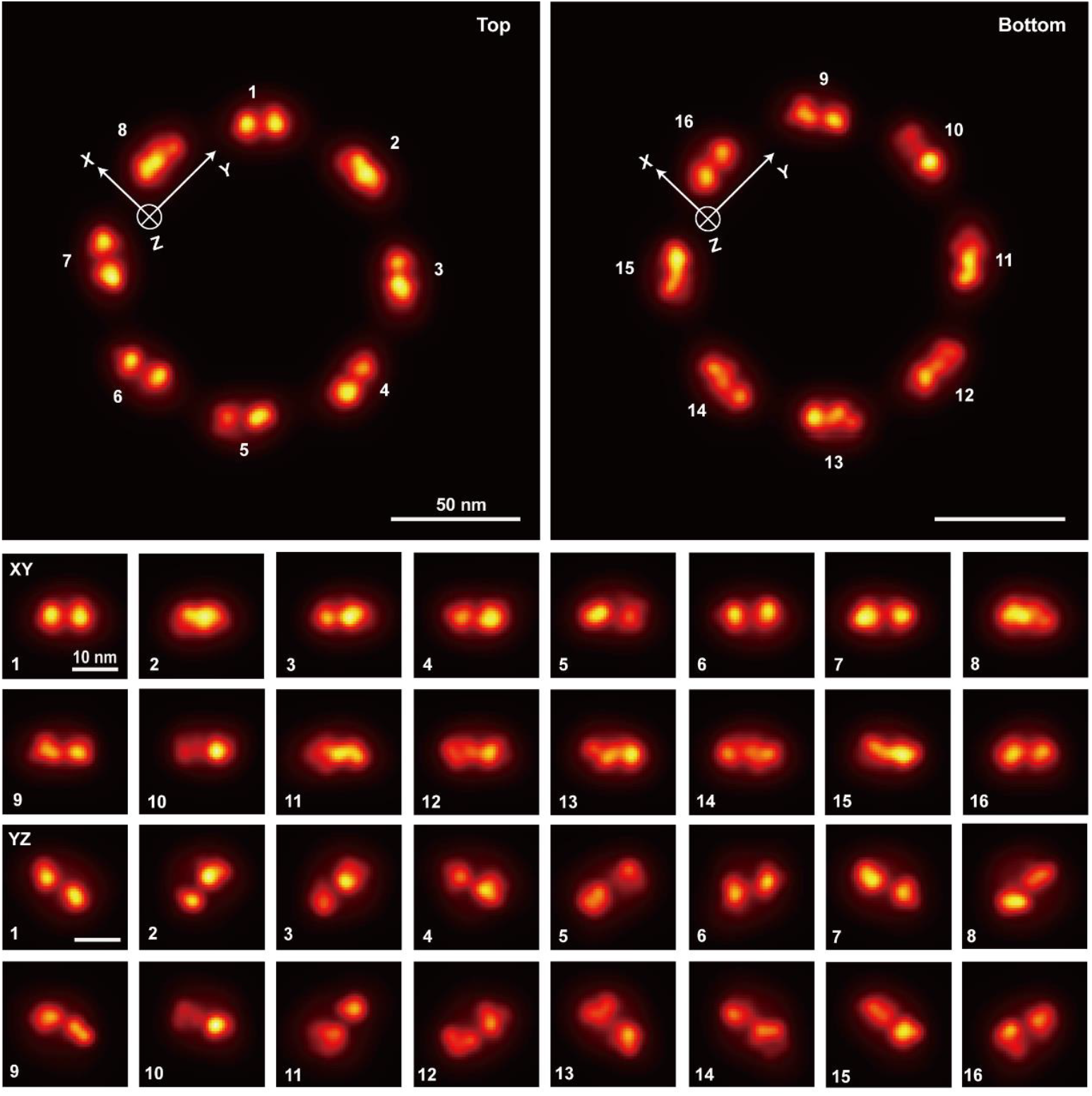
DeepSRFusion distinguished two tilted sub-modules within individual Nup96 subunits. The two upper panels present the three-dimensional projections of two ring structures, showing the spatial distribution of labeled sites (numbered 1-8, 9-16), with the XYZ coordinate system indicated, and the scale bars were 50 nm. The lower subpanels are divided into two imaging planes (XY and YZ), and each column corresponds to the local super-resolution imaging of different numbered sites on the rings, demonstrating the positional and morphological features of each labeled site in three-dimensional space and facilitating the analysis of the spatial organization rules of the ring structures. Bars were 50 nm in panels, 10 nm in subpanels.

**Extended Data Fig. 9:**
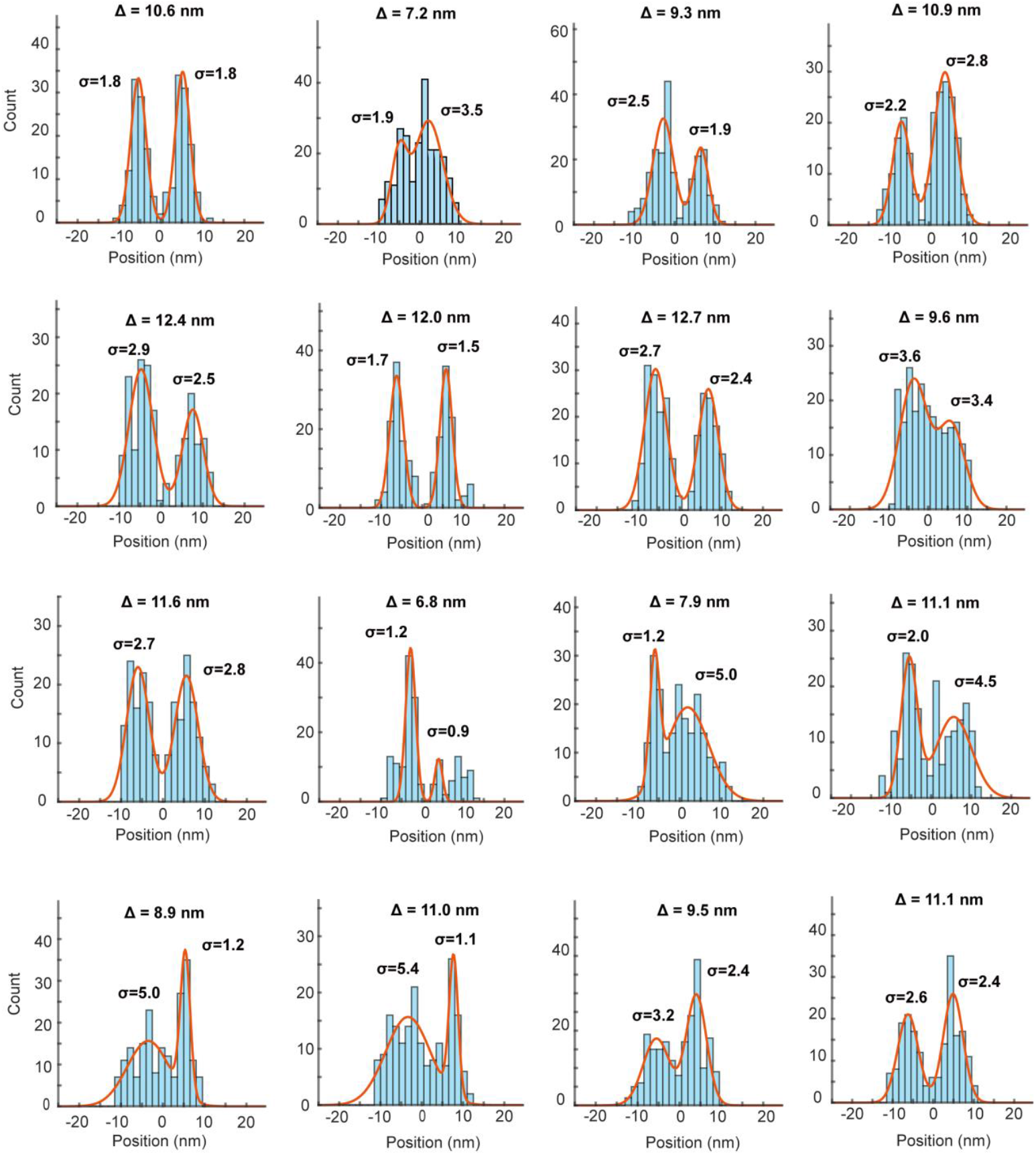
Histograms of two tilted sub-modules within individual Nup96 subunits. Histograms depicting the distribution of particle count corresponding to positions, with red curves representing fitted Gaussian profiles. Each subplot is annotated with parameters Δ characterizing spatial spacing and σ indicating the spread of the distribution, illustrating the spatial variability and fitting results of particle localization in super-resolution imaging analyses.

**Extended Data Fig. 10:**
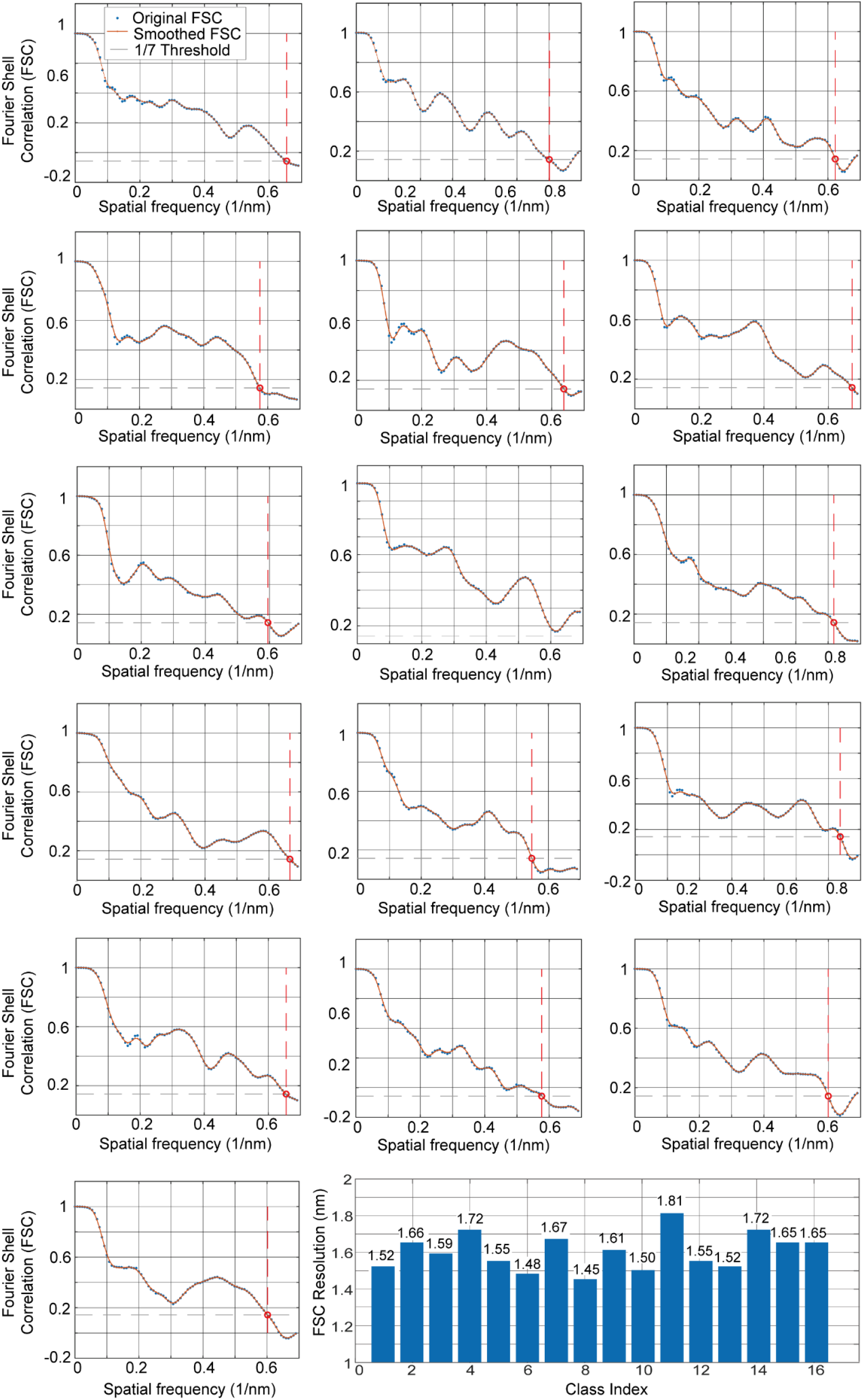
FSC analysis of the Nup96 subunits reconstructed by DeepSRFusion. FSC (**Supplementary Note 5.2**) curves for each subgroup were calculated independently, and the Fig. displays original FSC values (Original FSC), cubic-spline–smoothed FSC curves (Smoothed FSC), and the 1/7 resolution threshold (1/7 Threshold). The FSC resolution of each subgroup is defined as the inverse of the frequency at which the FSC curve first intersects the threshold. The bar plot at the bottom summarizes the FSC resolutions of all 16 subgroups, with an average resolution of 1.60 ± 0.10 nm, demonstrating that the reconstructed Nup96 achieves high-quality nanometer-scale structural resolution.

## Reference

[1] Sigal YM, Zhou R, Zhuang X. Visualizing and discovering cellular structures with super-resolution microscopy[J]. Science, 2018, 361: 880–887.

[2] Liu S, Hoess P, Ries J. Super-resolution microscopy for structural cell biology[J]. Annual Review of Biophysics, 2022, 51: 301–326.

[3] Lelek M, Gyparaki MT, Beliu G, et al. Single-molecule localization microscopy[J]. Nature Reviews Methods Primers, 2021, 1: 39.

[4] Sydor AM, Czymmek KJ, Puchner EM, Mennella V. Super-resolution microscopy: from single molecules to supramolecular assemblies[J]. Trends in Cell Biology, 2015, 25: 730–748.

[5] Brenlla A, Deen L, Annibale P. Single-molecule localization microscopy error is sensor dependent and larger than theory predicts[J]. Biophysical Reports, 2025, 5: 100223.

[6] Liu S, Huh H, Lee SH, Huang F. Three-dimensional single-molecule localization microscopy in whole-cell and tissue specimens[J]. Annual Review of Biomedical Engineering, 2020, 22: 155–184.

[7] Möckl L, Moerner WE. Super-resolution microscopy with single molecules in biology and beyond-essentials, current trends, and future challenges[J]. Journal of American Chemical Society, 2020, 142: 17828–17844.

[8] Sieben C, Douglass KM, Guichard P, Manley S. Super-resolution microscopy to decipher multi-molecular assemblies[J]. Current Opinion of Structural Biology, 2018, 49: 169–176.

[9] Kudryashev M, Castaño-Díez D, Stahlberg H. Limiting factors in single particle cryo electron tomography [J]. Computational and Structural Biotechnology Journal, 2012, 1: e201207002.

[10] Piper SJ, Johnson RM, Wootten D, Sexton PM. Membranes under the magnetic lens: a dive into the diverse world of membrane protein structures using cryo-EM[J]. Chemical Reviews, 2022, 122: 13989–14017.

[11] Mendes A, Heil HS, Coelho S, et al. Mapping molecular complexes with super-resolution microscopy and single-particle analysis[J]. Open Biology, 2022, 12: 220079.

[12] Salas D, Le Gall A, Fiche JB, et al. Angular reconstitution-based 3D reconstructions of nanomolecular structures from superresolution light-microscopy images[J]. Proceedings of the National Academy Sciences, 2017, 114: 9273–9278.

[13] Sieben C, Banterle N, Douglass KM, Gönczy P, Manley S. Multicolor single-particle reconstruction of protein complexes[J]. Nature Methods, 2018, 15: 777–780.

[14] Szymborska A, De Marco A, Daigle N, et al. Nuclear pore scaffold structure analyzed by superresolution microscopy and particle averaging[J]. Science, 2013, 341: 655–658.

[15] Liu J, Li Y, Chen T, Zhang F, Xu F. Machine learning for single-molecule localization microscopy: from data analysis to quantification[J]. Analytic Chemistry, 2024, 96: 11103–11114.

[16] Schnitzbauer J, Wang Y, Zhao S, et al. Correlation analysis framework for localization-based superresolution microscopy[J]. Proceeding of the National Academy of Science, 2018, 115: 3219–3224.

[17] Broeken J, Johnson H, Lidke DS, et al. Resolution improvement by 3D particle averaging in localization microscopy[J]. Methods and Applications in Fluorescence, 2015, 3: 014003.

[18] Vasan R, Ferrante AJ, Borensztejn A, et al. Interpretable representation learning for 3D multi-piece intracellular structures using point clouds[J]. Nature Methods, 2025, 22: 1531–1544.

[19] Levet F, Sibarita JB. PoCA: a software platform for point cloud data visualization and quantification[J]. Nature Methods, 2023, 20: 629–630.

[20] Heydarian H, Schueder F, Strauss MT, et al. Template-free 2D particle fusion in localization microscopy[J]. Nature Methods, 2018, 15: 781–784.

[21] Heydarian H, Joosten M, Przybylski A, et al. 3D particle averaging and detection of macromolecular symmetry in localization microscopy[J]. Nature Communications, 2021, 12: 2847.

[22] Wu YL, Hoess P, Tschanz A, et al. Maximum-likelihood model fitting for quantitative analysis of SMLM data[J]. Nature Methods, 2023, 20: 139–148.

[23] Hugelier S, Tang Q, Kim HH, et al. ECLiPSE: a versatile classification technique for structural and morphological analysis of 2D and 3D single-molecule localization microscopy data[J]. Nature Methods, 2024, 21: 1909–1915.

[24] Jordan MI, Mitchell TM. Machine learning: trends, perspectives, and prospects[J]. Science, 2015, 349: 255–260.

[25] Greener JG, Kandathil SM, Moffat L, Jones DT. A guide to machine learning for biologists[J]. Nature reviews Molecular Cell Biology, 2022, 23: 40–55.

[26] Yuan W, Eckart B, Kim K, et al. DeepGMR: learning latent gaussian mixture models for registration[C]. European Conference on Computer Vision, 2020: 733–750.

[27] Mei G, Poiesi F, Saltori C, et al. Overlap-guided gaussian mixture models for point cloud registration[C]. IEEE/CVF Winter Conference on Applications of Computer Vision (WACV), 2023: 4500–4509.

[28] Fu K, Luo J, Luo X, et al. Robust point cloud registration framework based on deep graph matching[J]. IEEE Transactions on Pattern Analysis and Machine Intelligence, 2022: 1–13.

[29] Nieves DJ, Pike JA, Levet F, et al. A framework for evaluating the performance of SMLM cluster analysis algorithms[J]. Nature Methods, 2023, 20: 259–267.

[30] Mazidi H, Ding T, Nehorai A, Lew MD. Quantifying accuracy and heterogeneity in single-molecule super-resolution microscopy[J]. Nature Communications, 2020, 11: 6353.

[31] Wu YL, Tschanz A, Krupnik L, Ries J. Quantitative data analysis in single-molecule localization microscopy[J]. Trends in Cell Biology, 2020, 30: 837–851.

[32] Thevathasan JV, Kahnwald M, Cieśliński K, et al. Nuclear pores as versatile reference standards for quantitative superresolution microscopy[J]. Nature Methods, 2019, 16: 1045–1053.

[33] Beck M, Hurt E. The nuclear pore complex: understanding its function through structural insight[J]. Nature Reviews Molecular Cell Biology, 2017, 18: 73–89.

[34] Walther TC, Alves A, Pickersgill H, et al. The conserved Nup107-160 complex is critical for nuclear pore complex assembly[J]. Cell, 2003,113: 195–206.

[35] Reinhardt S C M, Masullo L A, Baudrexel I, et al. Ångström-resolution fluorescence microscopy[J]. Nature, 2023, 617: 711–716.

[36] Sau A, Schnorrenberg S, Huang Z, et al. Overlapping nuclear import and export paths unveiled by two-colour MINFLUX[J]. Nature, 2025, 640: 821–827.

[37] Roy R, Al-Hashimi HM. AlphaFold3 takes a step toward decoding molecular behavior and biological computation[J]. Nature Structural & Molecular Biology, 2024, 31: 997–1000.

[38] Wang J, Tan C, Gao Z, et al. End-to-end cryo-EM complex structure determination with high accuracy and ultra-fast speed[J]. Nature Machine Intelligence, 2025, 7: 1091–1103.

[39] Tian Z, Gong J, Crowe M, et al. Biochemical studies of membrane fusion at the single-particle level[J]. Progress in Lipid Research, 2019, 73: 92–100.

[40] Klupp BG, Mettenleiter TC. The knowns and unknowns of herpesvirus nuclear egress[J]. Annual Review of Virology, 2023, 10: 305–323.

[41] Lancaster CE, Fountain A, Dayam RM, et al. Phagosome resolution regenerates lysosomes and maintains the degradative capacity in phagocytes[J]. Journal of Cell Biology, 2021, 220: e202005072.

[42] Bénac N, Ezequiel Saraceno G, Butler C, et al. Non-canonical interplay between glutamatergic NMDA and dopamine receptors shapes synaptogenesis[J]. Nature Communications, 2024, 15: 27.

